# Fucoidans are senotherapeutics that enhance SIRT6-dependent DNA repair

**DOI:** 10.1101/2025.04.27.650852

**Authors:** Lei Justan Zhang, Osama Elsallabi, Carolina Soto-Palma, Josh Bartz, Rahagir Salekeen, Allancer Nunes, Wandi Xu, Kyooa Lee, Brian Hughes, Borui Zhang, Abdalla Mohamed, Sara J. McGowan, Luise Angelini, Ryan O’Kelly, Seyed A. Biashad, Eric Hillpot, Francesco Morandini, Andrei Seluanov, Vera Gorbunova, Xiao Dong, Laura J. Niedernhofer, Paul D. Robbins

## Abstract

Aging is marked by the accumulation of senescent cells (SnCs), which contribute to tissue dysfunction and age-related diseases. Senotherapeutics, including senolytics which specifically induce lysis of SnCs and senomorphics, which suppress the senescence phenotype, represent promising therapeutic interventions for mitigating age-related pathologies and extending healthspan. Using a phenotypic-based senescent cell screening assay, we identified fucoidans, a class of sulfated polysaccharides derived from brown algae and seaweed, as novel senotherapeutics. In particular, fucoidan from *Fucus vesiculosus* (Fucoidan-FV) displayed potent senomorphic activity in different types of SnCs, reduced senescence in multiple tissues in aged mice, and extended healthspan in a mouse model of accelerated aging. Fucoidan-FV also enhanced the deacetylation and mono-ADP-ribosylation (mADPr) activity of SIRT6 and improved DNA repair and reduced senescence, in part, through SIRT6-dependent pathways. In addition, Fucoidan-FV downregulated genes associated with inflammation, Wnt signaling, and ECM remodeling pathways in SnCs and increased expression of genes involved with DNA repair. These findings support the translational potential of fucoidans as novel senotherapeutics that also are able to improve SIRT6-mediated DNA repair.

## INTRODUCTION

Aging is characterized by a progressive decline in physiological functions and overall fitness, driven by diverse mechanisms that result in accumulated cellular damage and impaired tissues homeostasis.^1–3^ One of the key hallmarks in this decline is cellular senescence, a state of cell cycle arrest triggered by external and internal stressors, such as DNA damage, oxidative stress, telomere shortening, or activation of oncogenes.^4,5^ These types of chronic stress result in the increased expression of cyclin-dependent kinase inhibitors, such as p16^INK4a^ and p21^Cip1^, which activate the retinoblastoma (pRB) and p53 pathways to halt cell proliferation. A common marker of senescent cells (SnCs) is senescence-associated β-galactosidase (SA-β-gal) activity.^6^ SnCs also develop a distinctive senescence-associated secretory phenotype (SASP), characterized by the secretion of pro-inflammatory cytokines, chemokines, metabolites, extracellular vesicles, and other factors that promote tissue remodeling and immune cell recruitment.^7^ While SASP components can support beneficial processes like wound healing, their chronic presence leads to sustained inflammation, impaired tissue regeneration, and functional decline. As SnCs accumulate with age, their SASP contributes to driving aging and the progression of numerous age-related diseases, including osteoarthritis, fibrosis, diabetes, cancer, and cardiovascular diseases.^8–10^

Numerous animal studies have demonstrated that reducing the SnC burden can improve physical function, reduce inflammation, and extend healthspan in animal models.^11–15^ Pharmacologic interventions targeting SnCs, termed senotherapeutics, have emerged as an effective approach to alleviate age-related pathology and extend healthspan, at least in animal models.^16,17^ Senotherapeutics can be broadly categorized as senolytics, which selectively eliminate SnCs, and senomorphics, which suppress the senescence phenotype, in particular the SASP, without killing SnCs.^18,19^ The discovery of natural compounds with senotherapeutic properties is particularly appealing due to their typically lower side effects and higher translational potential as safe interventions in aging.^20,21^

Fucoidans are a natural complex class of sulfated polysaccharides derived primarily from brown algae and seaweed. Structurally, fucoidans are unique in that they contain sulfate esters linked to specific carbon atoms on the (1→3) or alternating (1→3) and (1→4)-linked α-L-fucopyranose repeating units. Fucoidans have been found to exhibit a wide range of biological activities, functioning as anticoagulants, antioxidants, anti-inflammatories, antivirals, and with anti-cancer properties.^22–27^ Due to these diverse bioactivities, fucoidans have gained interest in the pharmaceutical, dietary supplement, and cosmetic industries as valuable natural compounds. However, whether the observed biological activities of fucoidans are mediated, in part, through a senotherapeutic activity is unknown. Here we screened a small library of fucoidans derived from various species of brown seaweed for senotherapeutic effect in several different types of SnCs. Although all fucoidans had some serotherapeutic activity, fucoidan derived from *Fucus vesiculosus* (Fucoidan-FV) emerged as the most potent senomorphic agent capable of reducing markers of tissue senescence and extending healthspan in mice whereas fucoidan from *Macrocystis pyrifera* (Fucoidan-MP) had senolytic activity. Interestingly, we and the accompanying paper by Biashad *et al*, showed that Fucoidan-FV also was the most effective fucoidan at stimulating the mono-ADP-ribosylation (mADPr) activity of SIRT6. We also demonstrate that Fucoidan-FV enhanced DNA repair, in part, in a SIRT6-dependent manner, reduced SASP expression and modulated multiple signaling pathways associated with cell cycle progression, canonical Wnt signaling, ECM remodeling, cell death regulation and DNA repair. Given the ability of Fucoidan-FV to also extend median lifespan in male mice, reduce frailty and epigenetic age in both sexes and repress expression of LINE1 elements, which requires the mADPr activity of SIRT6, taken together, these results strongly support the translational use of fucoidans to extend healthspan.

## RESULTS

### Phenotypic senescent cell-based screening identified senotherapeutic activity of Fucoidans

Fucoidans are complex, heterogeneous polysaccharides whose bioactivities vary significantly depending on differences in molecular structure including molecular weight, sulfation degree, and sugar composition. Their structural variations are influenced by multiple factors, such as seaweed species, geographic origin, harvesting season, harvest location, and extraction method.^28–30^ To capture their diversity in structures and bioactivities, we compiled a small library of fucoidans derived from diverse sources of brown seaweed that were commercially available.

To evaluate the senotherapeutic potential of fucoidans, we conducted a screen based on the phenotypic SA-β-gal activity of SnCs. Initially, primary *Ercc1*^-/-^ mouse embryonic fibroblasts (MEFs) were used as the SnCs model. These MEFs are highly susceptible to senescence under oxidative stress due to their deficiency in ERCC1, a component of the heterodimeric 3’ endonuclease with XPF involved in multiple types of DNA repair.^31^ This sensitivity makes *Ercc1*^-/-^ MEFs an ideal model for studying senescence-related cellular responses under stress conditions including oxidative stress induced by passage at 20% O_2_.^9,16,32^ Wild-type (WT) MEFs, which retain normal proliferative and DNA repair capacities, were used as non-senescent controls (NSnCs). Each type of fucoidan was tested in both *Ercc1*^-/-^ and WT MEFs over a 48-hour treatment period. A fluorogenic dye C_12_FDG then was used to stain for the SA-β-gal activity.^6,33^ Senescence levels were quantified based on the percentage of C_12_FDG-positive cells in each sample (**Fig. 1A**) using a high content fluorescent imager.

**Figure 1.**
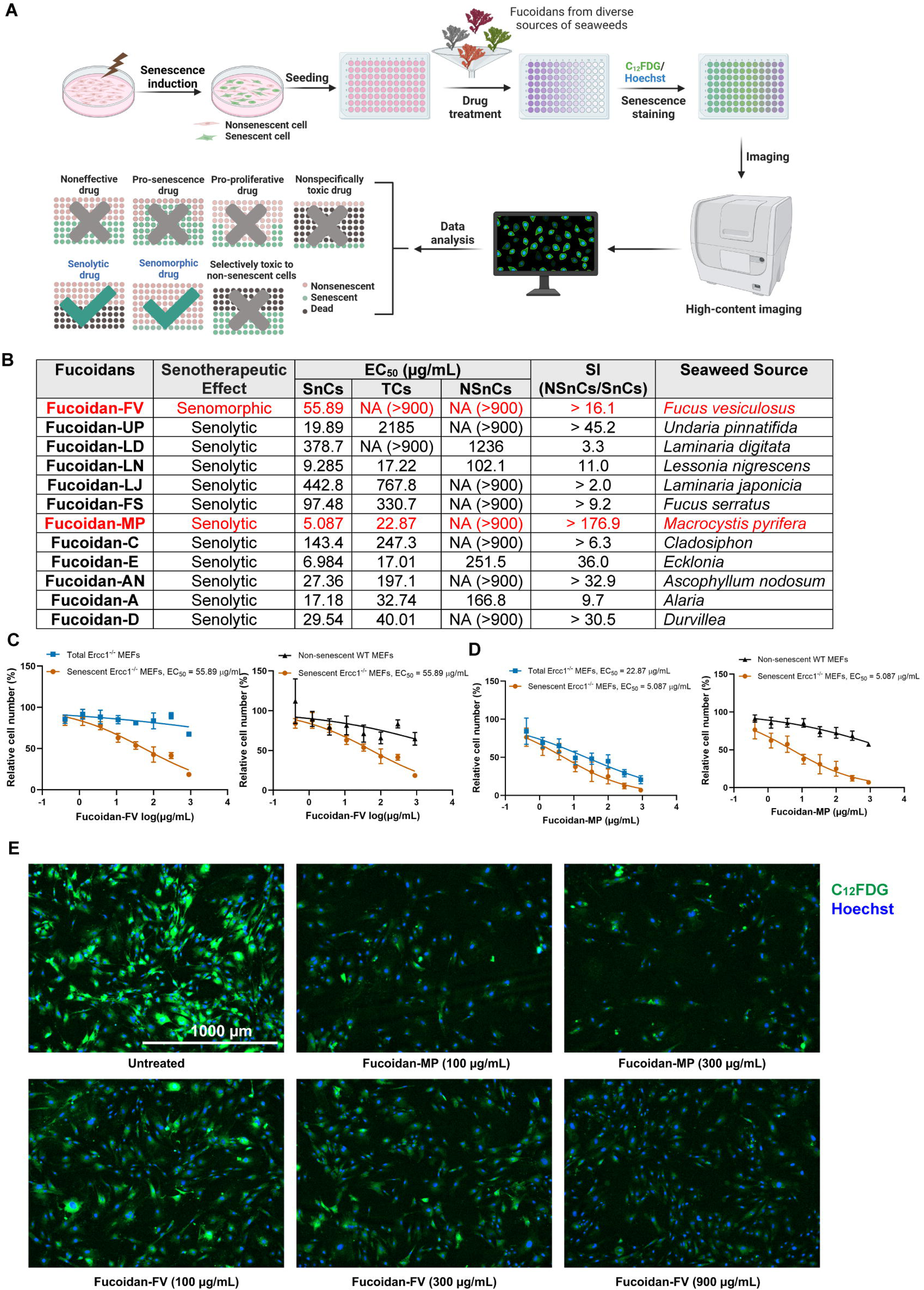
Identification of Fucoidans as a novel class of senotherapeutics through senescent cell-based phenotypic drug screening. (**A**) Focused library screening of different seaweed sources of Fucoidan in senescent *Ercc1*^-/Δ^ MEFs induced under oxidative stress. *Created in BioRender.com* (**B**) Summary of senescent cell screening of various Fucoidans in non-senescent WT MEF and senescent *Ercc1*^-/-^ MEF cells. Abbreviations: **SnCs**: C_12_FDG-positive senescent cells; **TCs**: Total *Ercc1*^-/-^ MEF cells; **NSnCs**: Non-senescent WT MEF cells; **SI**: Selectivity index. **EC_50_**: Concentration resulting in a 50% reduction in cell number compared to control. **NA**: EC_50_ values are not available due to inactivity. (**C**) Dose-response analysis of the senotherapeutic activities of Fucoidan-FV and Fucoidan-MP. Error bars represent SD for n = 3. (**D**) Representative images of the senotherapeutic effects of Fucoidan-FV and Fucoidan-MP on senescent *Ercc1*^-/-^ MEF cells. Images were captured using Cytation 1 at 4X magnification. Blue fluorescence indicates Hoechst 33324-stained nuclei and green fluorescence marks C_12_FDG-stained SA-β-gal positive senescent cells.

To distinguish between senolytic and senomorphic effects, we determined the effective concentration (EC_50_) for each fucoidan for both the number of C_12_FDG-positive *Ercc1*^-/-^ MEFs and the total cell number. Compounds that reduced both the total cell population and C_12_FDG-positive cells were classified as potential senolytics, while those that reduced C_12_FDG-positive cells without affecting total cell count were categorized as senomorphics. The selectivity index (SI), calculated as the EC_50_ ratio between NSnCs and SnCs, provided a metric to assess the selectivity of the compounds.

All of the tested fucoidan variants exhibited ability to reduce the number of SA-βgal+ cells, consistent with senotherapeutic activity, though their potency and selectivity varied (**Fig. 1B**). Notably, fucoidan from *Fucus vesiculosus* (Fucoidan-FV) displayed the strongest senomorphic effect, effectively reducing the percent of SA-β-gal+ cells associated with the senescent phenotype without inducing cell death. In contrast, fucoidan from *Macrocystis pyrifera* (Fucoidan-MP) showed a pronounced senolytic effect, significantly reducing the senescent cell population by inducing SnC death specifically (**Fig. 1C-E**). These results suggest that fucoidans from different sources exhibit distinct senotherapeutic activities, similar to their diverse bioactivities previously observed.

To further confirm the senotherapeutic activities, we extended our analysis to additional cell types and inducers of senescence. These included WT MEF cells induced to senescence through oxidative stress (H_2_O_2_) and genotoxic stress (etoposide), human IMR90 fibroblast cells exposed to genotoxic agent etoposide, and replicative senescent human umbilical vein endothelial cells (HUVECs) induced through repeated passaging. Across these diverse senescent cell models, Fucoidan-FV consistently demonstrated the strongest senomorphic effect by reducing the number of C_12_FDG-positive cells (**Fig. S1**), whereas Fucoidan-MP displayed senolytic activity by eliminating SnCs specifically (**Fig. S2**). Based on these results, we selected the senomorphic Fucoidan-FV and the senolytic Fucoidan-MP for testing *in vivo*.

### Senotherapeutic effects in naturally aged mice

To evaluate the senotherapeutic effects of Fucoidan-FV and Fucoidan-MP *in vivo*, we administered these compounds by oral gavage to naturally aged female WT C57BL/6 mice (32 months old). The mice received daily doses of either 1 g/kg of Fucoidan-FV or 500 mg/kg of Fucoidan-MP for five consecutive days. Two days after the last dose, tissues were collected for analysis (**Fig. 2A and 2E**). Molecular analysis demonstrated that both fucoidans reduced markers of cellular senescence and SASP in multiple tissues (**Fig. 2B and 2F**), with particularly effects in the kidney (**Fig. 2C**) and lung, tissues known to accumulate high levels of SnCs with aging. Additionally, both fucoidan treatments reduced SASP markers in quadriceps muscles, such as genes encoding TNF-α, IL-6, IL-1β, CXCL1, and MCP1 (**Fig. 2B, 2F, and S3**). Notably, Fucoidan-FV treatment also led to a significant reduction in p21^Cip1^-positive γδ T cells in the spleen (**Fig. 2D**), a distinct subset of T lymphocytes involved in tissue homeostasis, immune surveillance, and early stress responses. Tissue-resident γδ T cells accumulate with age, particularly in visceral adipose tissue, where they contribute to chronic low-grade inflammation and metabolic dysfunction.^34^ This observed reduction of senescent γδ T cells suggests a potential senotherapeutic effect of fucoidan on specific immune cell populations in aged mice. Overall, Fucoidan-FV demonstrated greater senotherapeutic effects than Fucoidan-MP in these naturally aged mice, at least at the dosing regimen used.

**Figure 2.**
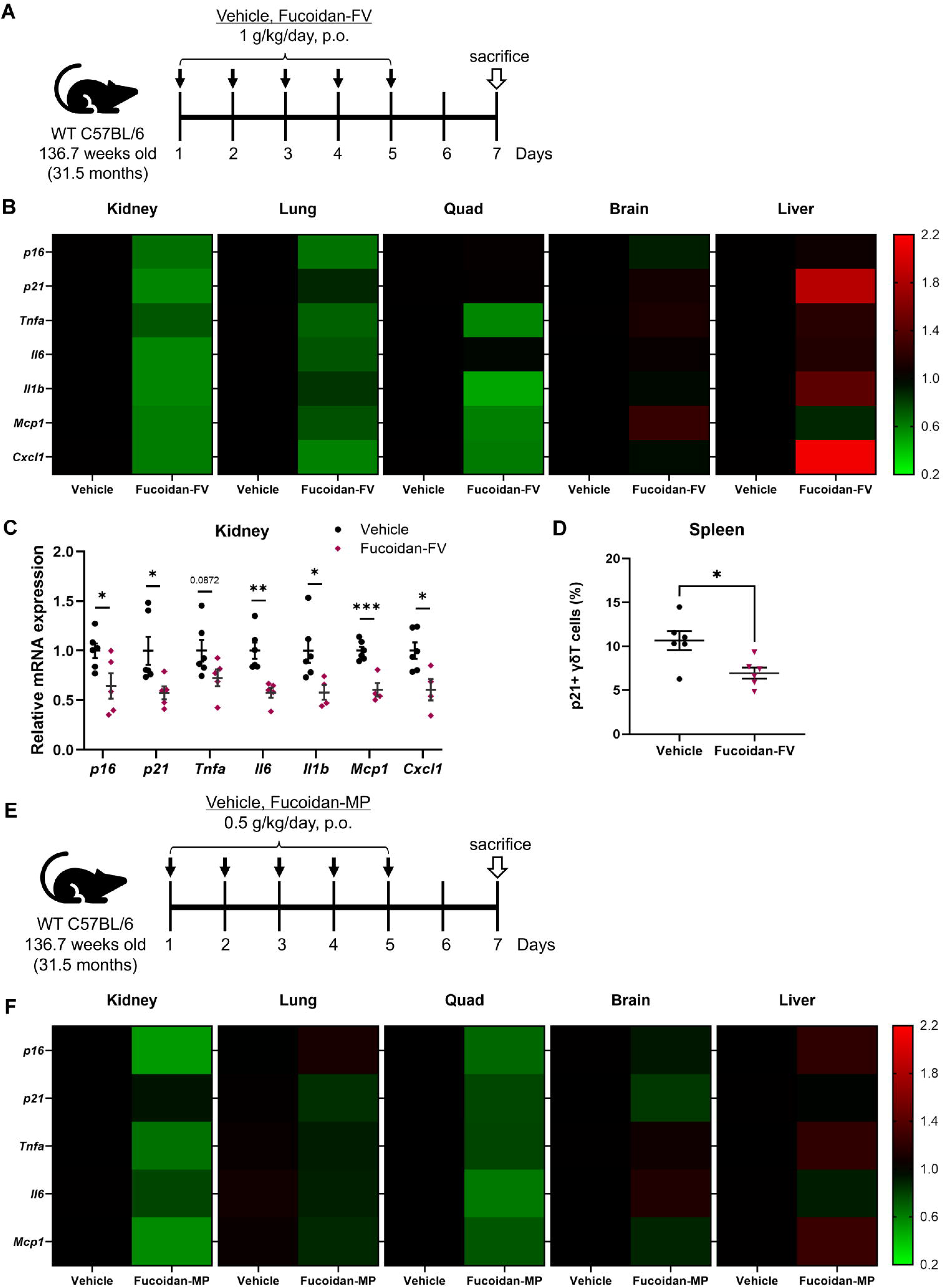
Evaluation of the senotherapeutic effects of Fucoidan-FV and Fucoidan-MP in naturally aged mice. (**A**) WT C57BL/6 mice aged 32 months old were treated with 1 g/kg of Fucoidan-FV by oral gavage for five consecutive days. Tissues were collected two days after the last dose for analysis. (**B**) Effects of Fucoidan-FV on reducing senescence across different tissues in aged WT C57BL/6 mice, especially in the (**C**) kidney. Error bars represent SEM for n = 6. (**D**) Fucoidan-FV reduced p21+ senescent γδ T cells in the spleen of the 32-month-old WT C57BL mice. Error bars represent SEM for n = 6. (**E**) WT C57BL/6 mice aged 32 months old were treated with 1 g/kg of Fucoidan-MP by oral gavage for five consecutive days. Tissues were collected two days after the last dose for analysis. (**F**) Effects of Fucoidan-MP on reducing senescence across different tissues in aged WT C57BL/6 mice.

### Fucoidan-FV extends healthspan in progeroid mice

We then tested the ability of Fucoidan-FV to extend healthspan in the *Ercc1*^-/Δ^ progeria mouse model of accelerated senescence and aging.^35^ The *Ercc1*^-/Δ^ mice have reduced ERCC1 expression and thus higher sensitivity to DNA damage, increased accumulation of SnCs, premature aging phenotypes, and 6 fold shorter lifespan.^31^ Fucoidan-FV was incorporated into the diet at a concentration of 5% and administered ad libitum to male and female *Ercc1*^-/Δ^ mice over a six week period starting from 10 weeks of age (**Fig. 3A**). Weekly health assessments were conducted to monitor age-related symptoms, including tremor, kyphosis, dystonia, ataxia, gait disorder, hindlimb paralysis, and forelimb grip strength. Fucoidan-FV effectively reduced the composite score of aging symptoms, particularly during weeks 11 to 12 (**Fig. 3B**), an age where many aging symptoms first appear. Further analysis using RT-qPCR revealed a notable reduction in senescence markers and SASP factors, especially in the kidney and lung tissues (**Fig. 3C and 3D**). In contrast, the effects of Fucoidan-FV on tissue senescence were less pronounced in the brain and liver (**Fig. 2B, 2F, and 3C**). Together, these results confirm that Fucoidan-FV exhibits promising senotherapeutic effects in mouse models of natural and accelerated aging.

**Figure 3.**
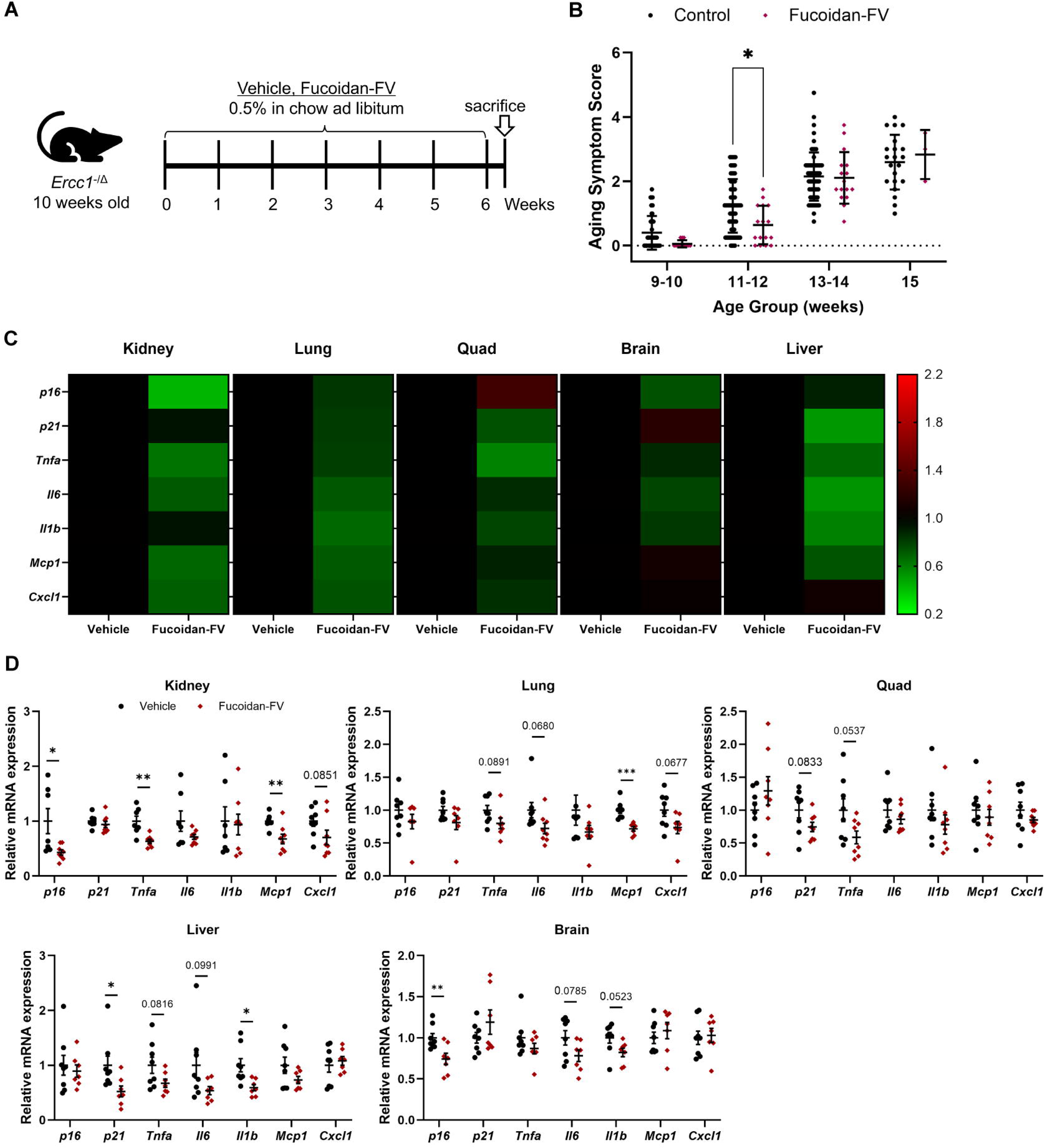
Evaluation of the senotherapeutic effects of Fucoidan-FV in accelerated aging mice. (**A**) *Ercc1*^-/Δ^ progeria mice at 10 weeks old were treated with 0.5% Fucoidan-FV in chow ad libitum for six consecutive weeks. Mice were sacrificed two days after the end of the treatment. (**B**) Fucoidan-FV treatment extended healthspan *Ercc1*^-/Δ^ progeria mice. Weekly health assessments were conducted to score age-related symptoms, including tremor, kyphosis, dystonia, ataxia, gait disorder, hindlimb paralysis, and forelimb grip strength. (**C**) Chronic Fucoidan-FV treatment reduced senescence in multiple tissues of *Ercc1*^-/Δ^ progeria mice, (**D**) including kidney, lung, quadriceps, liver, and brain. Error bars represent SEM for n = 8.

### Fucoidan treatment results in transcriptomic downregulation of SASP genes and increased expression of DNA repair processes

In order to elucidate potential molecular mechanisms underlying the observed senotherapeutic properties of Fucoidan, bulk RNA sequencing (RNA-Seq) was performed on non-senescent (control) and senescent MEFs treated with Fucoidan-FV. WT MEFs were treated with etoposide to induce senescence or vehicle to act as a control. The etoposide induced SnCs were then treated with Fucoidan-FV at 100 µg/mL or 300 µg/mL doses and subjected to bulk RNA-Seq analysis. The RNA-seq data confirmed that the SnCs indeed have a senescent transcriptome (**Fig. S4A**) with significant upregulation of CDKN1A and CDKN2A (**Fig. S4B**) as well as a variety of SASP factors found in the SenMayo gene panel^36^ (**Fig. 4A and S4C**). To identify transcriptional changes conferred by Fucoidan-FV treatment, differential expression analysis was performed using DESeq2. Although cell cycle arrest markers like CDKN1A (p21^Cip1^) and CDKN2A (p16^Ink4a^) were unchanged, Fucoidan-FV treatment reduced the expression of a variety of SASP factors (**Fig. 4A**), supporting the senomorphic activity of Fucoidan-FV.

**Figure 4.**
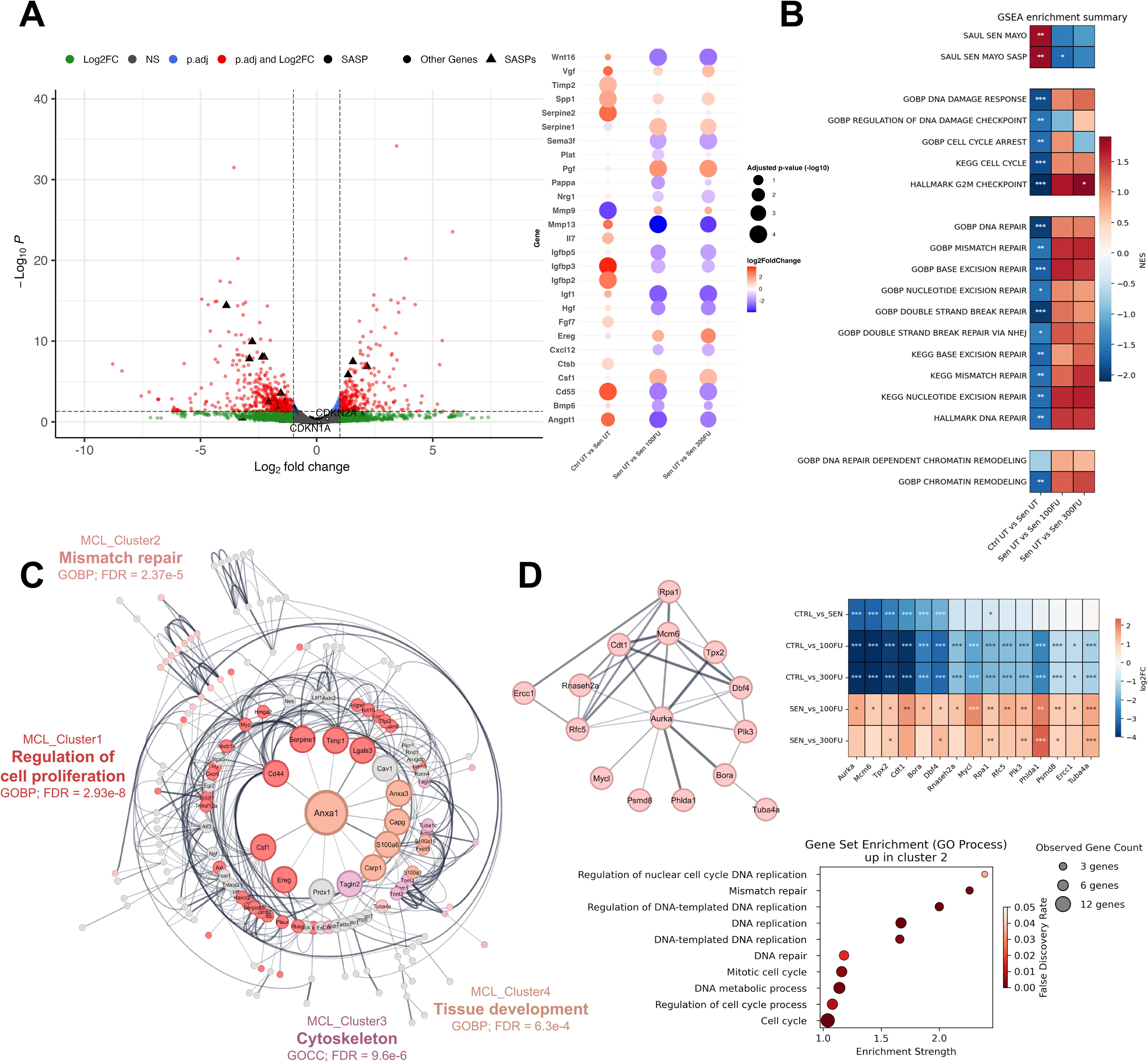
Bulk RNA-Seq analysis of transcriptomic changes in senescent MEFs treated with Fucoidan-FV. (**A**) Volcano Plot of DEGs between UT Senescent vs 100 µg/mL Fucoidan-FV treated MEFs (left). SASP genes are highlighted as enlarged black triangles, SASPS with significant changes are further elucidated in a bubble plot (right). (**B**) Normalized Gene set enrichment analysis (GSEA) scores for selected gene sets associated with senescence/SASPs, DNA damage response, and DNA repair processes in different treatment pairs. (**C**) Radial interaction network diagram of filtered upregulated DEGs in UT Sen vs 100 µg/mL Fucoidan-FV treated MEFs. Node size corresponds to degree centrality in network topology; edge width corresponds to undirected confidence of interaction between nodes. Top 4 MCL clusters are annotated with colored nodes and labeled with the highest enriched GO term. (**D**) Subnetwork of MCL cluster 2 gene interactions from panel C (left), expression changes of cluster 2 genes in different experimental group pairs (right), and enrichment of GO biological processes by cluster 2 genes (bottom).

To further elucidate the transcriptional effects of Fucoidan-FV and identify a potential mechanism(s) of action, we performed gene set enrichment analyses (GSEA). Fucoidan-FV treatment resulted in a negative enrichment of the Sen Mayo gene panel, particularly SASP genes (**Fig. 4B**) and a positive enrichment in gene sets associated with the cell cycle, the DNA damage response and the G2M checkpoint. Additionally, we interrogated multiple gene sets associated with DNA repair and found Fucoidan-FV-treated SnCs to have increased expression in genes in DSB repair, excision repair processes, and DNA repair dependent chromatin remodeling pathways, suggesting that Fucoidan-FV also enhanced overall DNA repair activity.

In order to identify potential Fucoidan-FV targets exclusive to SnCs, we constructed an interaction network of genes that were differentially up- or downregulated in MEFs treated with Fucoidan-FV compared to senescent MEFs, but not control cells (**Fig. 4C and S4D**). The constructed downregulated gene network contained many cytokines and SASP as degree central nodes including HGF, MMP13, DCN, CXCL5/15, and TGFBR2 and major clusters of these nodes associated with processes such as ECM remodeling, cellular migration, and canonical Wnt signaling (**Fig. S4E and S4F**). In the upregulated gene network, the central nodes included ANXA1/3, SERPINE1, CD44, CSRP1, and TIMP1. The majority of these genes were assigned to two major clusters primarily associated with cell proliferation and DNA/mismatch repair (**Fig. 4D and S4F**). Upregulated cluster 2 overrepresents multiple DNA repair associated processes and includes 15 major cell cycle and DNA repair machinery genes including ERCC1, RPA1, RFC5, MCM6, AURKA, and PLK3 (**Fig. 4E**). Interestingly, most of these genes were downregulated in untreated SnCs compared to control cells, indicating Fucoidan-FV mediates at least a partial reversal of these processes in SnCs. Taken together, these findings suggest fucoidan treatment results in upregulation of DNA repair processes in SnCs, which potentially results in its senomorphic activities including suppression of SASPs, reduced expression of ECM remodeling proteins, and positive regulation of cell cycle progression and proliferation.

### Fucoidan mediated DNA repair and SASP regulation is dependent on SIRT6 activity

It has been previously reported that fucoidans, particularly those from *Fucus distichus* and *Fucus vesiculosus*, enhanced Sirtuin 6 (SIRT6) deacetylase activity,^37^ an NAD+-dependent enzyme crucial for maintaining genomic stability, cellular stress responses, and longevity.^38–40^ Furthermore, in addition to the role of SIRT6 in PARP1-mediated DNA repair, it is also epigenetically regulates the expression of many cytokines as well as Wnt target genes. Given that we have demonstrated previously that a rare variant in SIRT6 found in centenarians had increased mono-ADP-ribosylation and DNA repair activities, we hypothesized that the transcriptomic alterations observed in our RNA-Seq analysis from Fucoidan-FV treatment is mediated by activation of SIRT6.

Since there were no significant transcriptomic changes in SIRT6 expression in our RNA-Seq analysis, we examined whether there were changes in SIRT6 enzymatic activity. SIRT6 has multiple enzymatic activities, including deacetylation of histone H3 lysine 9 (H3K9) and lysine 56 (H3K56),^41,42^ as well as mono-ADP-ribosylation.^43^ To explore the molecular mechanisms underlying the senotherapeutic effects of fucoidan-FV, we examined the effect on deacetylation of histone H3 lysine and self mono-ADP-ribosylation activity. Interestingly, Fucoidan-FV increased the deacetylation activity (**Fig. 5A**) and, to a greater extent, mADPr activity of SIRT6 (**Fig. 5B**).

**Figure 5.**
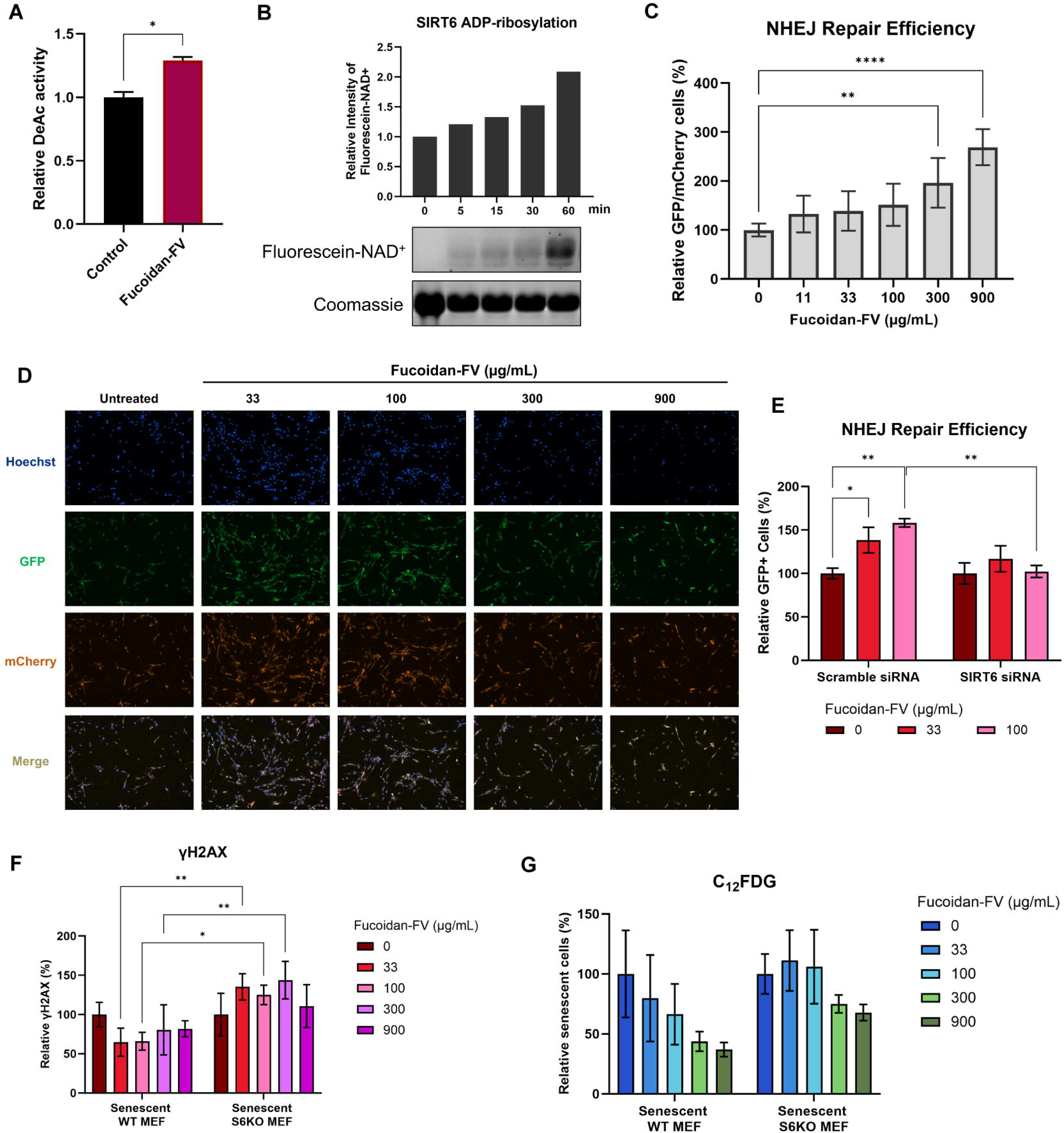
Fucoidan-FV induced senotherapeutic effects are mediated by SIRT6. (**A**) Fucoidan-FV (500 µg/mL) increases deacetylation activity of SIRT6. (**B**) Fucoidan-FV enhanced self-ribosylation of SIRT6 using fluorescein-labeled NAD+. Recombinant SIRT6 was incubated with NAD+ conjugated with a biotin residue and then run on an SDS-PAGE. (**C**) Fucoidan-FV enhances NHEJ-based DNA damage repair in a dose-response manner. Error bars represent SD for n = 3. (**D**) Representative images showing Fucoidan-FV increased the NHEJ repair efficiency in NHEJ-GFP reporter cells. Successful NHEJ repair and transfection efficiency were detected by GFP and mCherry signals, respectively. (**E**) NHEJ repair efficiency of Fucoidan-FV is partially dependent on SIRT6. MEF cells were transfected with control siRNA and SIRT6 siRNA. After siRNA transfection, cells were treated with Fucoidan-FV for 48 h. Error bars represent SD for n = 2. (**F**) The reduction of DNA damage by Fucoidan-FV is partially dependent on SIRT6 in MEF cells. Error bars represent SD for n = 2. (**G**) The senomorphic effect of Fucoidan-FV is partially dependent on SIRT6 in MEF cells. Error bars represent SD for n = 2. Data are expressed as mean ± SEM. *p < 0.05 compared with vehicle.

Given that the mono-ADP-ribosylation activity of SIRT6 is crucial for DNA damage repair, particularly through the non-homologous end-joining (NHEJ) pathway, a key mechanism for repairing DNA double-strand breaks,^43^ we examined whether Fucoidan-FV could increase NHEJ-mediated DNA repair. Using human fibroblast cells harboring an NHEJ-GFP reporter, the cells were co-transfected with a plasmid encoding an I-SceI endonuclease to induce double-strand breaks and an mCherry plasmid to monitor transfection efficiency. In this assay, successful DNA repair restores GFP expression following I-Sce1 cleavage in mCherry positive cells.^43^ Treatment with Fucoidan-FV led to a dose-dependent increase in GFP-positive cells, indicating enhanced NHEJ-mediated DNA repair capacities (**Fig. 5C and 5D**). To confirm that the observed DNA repair enhancement was dependent on SIRT6, an siRNA was used to knock down SIRT6 expression. While Fucoidan-FV significantly increased NHEJ repair in cells treated with control siRNA, this effect was abrogated in cells with SIRT6 knockdown (**Fig. 5E**). This dependence on SIRT6 was further validated in WT and SIRT6 knockout (S6KO) MEF cells, where Fucoidan-FV treatment reduced γH2AX-positive foci, a marker of DNA damage, in WT MEFs, but showed no effect in S6KO cells (**Fig. 5F**). These findings suggest that the DNA repair activity of Fucoidan-FV is mediated, at least in part, through SIRT6 activation.

We next investigated whether the senomorphic effects of Fucoidan-FV are also dependent on SIRT6. In etoposide-induced senescent WT MEFs, treatment with Fucoidan-FV led to a reduction in C_12_FDG-positive cells. However, this effect was significantly diminished in S6KO cells, suggesting that the senomorphic activity of Fucoidan-FV may partially contribute to SIRT6-mediated enhancement of DNA repair (**Fig. 5G**). Taken together, these results suggest that Fucoidan-FV mechanistically functions as an activator of SIRT6 to improved DNA repair capacity and suppression of senescence markers and SASP in SnCs.

## DISCUSSION

Unlike synthetic senotherapeutics, natural compounds derived from marine algae and seaweed are particularly attractive for therapeutic development because of their lower risk of side effects and their long-standing presence in the diets of many populations. Despite this, there have been limited studies to identify potential national products with senotherapeutic activity, the one exception being flavonoids such as fisetin, luteolin and quercetin. Given that fucoidans have been reported to have numerous biological activity that could provide a health benefit as well as activate enzymatic activity of SIRT6, here we examined the potential serotherapeutic activity of fucoidans from different types of brown seaweed.^44,45^ Using initially a senescent cell phenotype-based screening assay in ERCC1-deficient MEFs followed by additional assays, we demonstrated that fucoidans, particularly Fucoidan-FV from *Fucus vesiculosus*, are a novel class of senotherapeutics. Interestingly, although almost all the fucoidans tested had senomorphic activity, a related fucoidan from *Macrocystis pyrifera* (Fucoidan-MP) had a senolytic activity. Why this one type fucoidan was able to induce SnC death as compared to the other fucoidans is still unclear.

SIRT6 is a NAD+-dependent histone deacylase and deacetylase which regulates key cellular processes, including glucose metabolism, oxidative stress, genomic integrity, and longevity.^38–40^ A key mechanistic finding of this and the accompanying study is the activation of SIRT6 by Fucoidan-FV, which appears to enhance SIRT6 deacetylase and, in particular, mono-ADP-ribosylation activities, both essential for DNA repair and genomic stability.^46,47^ Specifically, we observed enhanced non-homologous end joining (NHEJ), a crucial DNA repair pathway under cellular stress, through SIRT6 activation. This finding aligns with previous studies showing that SIRT6 facilitates DNA repair by recruiting essential factors, such as 53BP1 and NBS1, to damaged sites, which requires both its deacetylase and ADP-ribosyltransferase activities.^43^ SIRT6 is also a major epigenetic regulator of many inflammatory cytokine expression including IL1B, IL2/6/10/13/, TNF-α, MCP-1 and other NF-κB target genes. In addition, the treatment of

SnCs with fucoidan also increased the expression of certain DNA repair genes, suggesting that fucoidan can increase SIRT6 dependent DNA repair activity through multiple pathways. These results support a model where SIRT6 mediated chromatin remodeling can contribute to suppression of many of these SASP genes in Fucoidan-FV treated cells. Additionally, SIRT6 mediated regulation of many stemness-associated processes, including canonical Wnt signaling and cell migration, could also contribute to the therapeutic effects conferred by Fucoidan-FV treatment in the different mouse models of aging. It is also likely that improving DNA repair directly contributes to the senomorphic activity of Fucoidan-FV.

In the accompanying paper by Biashad *et al*, Fucoidan-FV provided in the food resulted in a significant increase in median lifespan in male mice and a marked reduction in frailty and epigenetic age in both male and female mice. Fucoidan-FV also repressed expression of LINE1 elements, which is consistent with reduction in inflammation.^48^ Here it is important to note that the mADPr activity of SIRT6 is required for LINE1 inhibition, consistent with the ability of Fucoidan-FV to stimulate SIRT6 activity *in vivo*. Whether the improvement in the epigenetic clock in Fucoidan-FV treatment mice is due to increase SIRT6 deacetylase activity is unclear.

Although we have observed effects of fucoidans on SIRT6 activity *in vitro* and on SIRT6-dependent DNA repair *in vivo*, structurally, it is unlikely that a bulky sulfated fucan polysaccharide can infiltrate and interact with nuclear SIRT6. Intact fucoidans may not directly enter the nucleus, but still could potentially activate SIRT6 through upstream signaling pathways. In addition, it is possible that certain metabolic intermediates or breakdown products of fucoidan are responsible for the observed effects. *In vivo*, digestive and gut microbial components likely contribute to fucoidan metabolism, potentially resulting in more active metabolites.^49^ Identification of the components/metabolites of fucoidan that are responsible for conferring the beneficial effects on SIRT6 activation, DNA repair, senescence, inflammation, the epigenetic clock and healthspan should result in more effective fucoidan-based gerotherapeutics. Finally, it is important to note that the senotherapeutic effects of fucoidan in culture appear to be only partially dependent on SIRT6, especially at high concentrations. Given that fucoidan has been reported to bind to other targets including TLR4, there may be additional mechanisms through which fucoidan functions to extend healthspan and reduce inflammation.

In summary, here and in the accompanying paper, we used a high-content senescent cell-based phenotypic screen to identify fucoidans as a new class of natural senotherapeutics with distinct senomorphic and senolytic properties. Among them, *Fucus vesiculosus*-derived fucoidan (Fucoidan-FV) had the strongest senomorphic activity as well as strongest ability to stimulate the mono-ribosylation activity of SIRT6. Mechanistically, Fucoidan-FV restored expression of DNA repair genes and enhanced SIRT6 activity, promoting genomic stability and suppressed inflammatory phenotypes in SnCs. Treatment with Fucoidan-FV extended healthspan in aged mice, improving physical function and reducing tissue senescence burden without adverse effects. In the accompanying paper we demonstrate that fucoidan-FV extends lifespan and healthspan in wild type aged mice and silences LINE1 retrotransposable elements. Together, these findings reinforce the clinical potential of Fucoidans as promising safe, effective, and novel senotherapeutic agents. Importantly, given the established safety profile and biocompatibility, fucoidans from *Undaria pinnatifida* and *Fucus vesiculosus* have also been granted GRAS (Generally Recognized as Safe) status for their use as food ingredients by the FDA.^50^ This allows for broader exploration of fucoidans in both pharmaceutical and nutraceutical applications for reducing DNA damage, senescence and inflammation while extending healthspan.

## ACKNOWLEDGEMENTS

Funding for this research was provided by NIH grants R01 AG069819 (PDR), P01 AG043376 (PDR, LJN), U19 AG056278 (PDR, LJN, VG), RO1 AG063543 (LJN), P01 AG062413 (LJN, PDR), U54 AG079754 (LJN, PDR), U54 AG076041 (LJN, PDR) and T32AG029796 (CSP). P01 AG047200 (AS, VG), P01 AG051449 (AS, VG), R01 AG027237 (VG), R37 AG046320 (AS).

## AUTHOR CONTRIBUTIONS

LJZ, LJN, VG, AS and PDR designed experiments and interpreted the results. LJZ, OE, WX, KL, BH, BZ, AM, SJM, LA, SAB, EH, FM and RO performed experiments. JB, RS, and CSP performed bioinformatic and computational analyses. LJZ, JB, RS, CSP, and PDR prepared the original draft of the manuscript. All authors reviewed the manuscript. LJZ, LJN, AS, VG and PDR supervised research. PDR, LJN, AS and VG secured funding.

## METHODS

### Resource availability Lead contact

Further information and requests for resources and reagents should be directed to and will be fulfilled by the lead contact, Paul D. Robbins.

### Compounds and reagents

Fucoidans were purchased from Biosynth (Staad, Switzerland). Fucoidan-FV was purchased from Sigma-Aldrich (F8190). Fucoidan-MP was purchased from Sigma-Aldrich (F8065). Hoechst 33342 was purchased from ThermoFisher (H1399). C_12_FDG was purchased from Setareh Biotech (7188). Formaldehyde 32% was purchased from Electron Microscopy Sciences (15714).

### Cells and mice

Primary *Ercc1^-/-^* mouse embryonic fibroblasts (MEFs) and WT MEFs were isolated on embryonic day 12.5-13.5. In brief, mouse embryos were isolated from yolk sac followed by the complete removal of viscera, lung and heart if presented. Embryos were then minced into fine chunks, fed with MEFs medium, cultivated under 3% oxygen to reduce stress. Cells were split at 1:3 when reaching confluence. MEFs were grown at a 1:1 ratio of Dulbecco’s Modification of Eagles Medium (supplemented with 4.5 g/L glucose and L-glutamine) and Ham’s F10 medium, supplemented with 10% fetal bovine serum, penicillin, streptomycin and non-essential amino acid. To induce oxidative stress-mediated DNA damage, *Ercc1^-/-^* MEFs were switched to 20% oxygen for three passages. WT MEFs were induced to undergo senescence by treating them with hydrogen peroxide H_2_O_2_ (200 μM) or etoposide (2 μM) for 24 h, followed by 5 days in normal culture media.

Human IMR90 lung fibroblasts were obtained from American Type Culture Collection (ATCC) and cultured in EMEM medium with 10% FBS and pen/strep antibiotics. To induce senescence, cells were treated with 20 μM etoposide for 24 h, followed by five days in normal culture media.

Human umbilical vein endothelial cells (HUVECs) were obtained from ATCC and cultured using Endothelial Cell Growth Media plus supplement (without vascular endothelial growth factor (VEGF)) and 1% pen/strep antibiotics. The cells were experimentally treated at late passages 13 to 15.

*Ercc1*^+/−^ and *Ercc1*^+/Δ^ mice from C57BL/6J and FVB/n backgrounds were crossed to generate *Ercc1*^−/Δ^ mice to prevent potential strain-specific pathology. Aged wild-type C57BL/6J:FVB/NJ mice were generated by crossing C57BL/6J and FVB/n inbred mice purchased from Jackson Laboratory. Mice were left to age for two years before being enrolled into the late life intervention study. Animal protocols used in this study were approved by the University of Minnesota Institutional Animal Care and Use Committees.

### Senotherapeutic screening

Senescence was evaluated based on SA-β-gal activity using C_12_FDG staining assay. Specifically, senescent *Ercc1^−/−^* MEFs were passaged for three times at 20% O_2_ to induce senescence then seeded at 3000 cells per well in black wall, clear bottom 96 well plates at least 16 hours prior to treatment. Following the addition of drugs, the MEFs were incubated for 48 hours at 20% O_2_. After removing the medium, cells were incubated in 100 nM Bafilomycin A1 in culture medium for 60 min to induce lysosomal alkalinization, followed by incubation with 20 μM fluorogenic substrate C_12_FDG (7188, Setareh Biotech, USA) for 2 h and counterstaining with 2 μg/ml Hoechst 33342 (H1399, Thermo Fisher Scientific, MA, USA) for 15 min. Subsequently, cells were washed with PBS and fixed in 2% paraformaldehyde for 15 min. Finally, cells were imaged with 6 fields per well using a high content fluorescent image acquisition and analysis platform Cytation 1 (BioTek, VT, USA).

### Healthspan evaluation of *Ercc1^-/^*^Δ^ mice

Healthspan assessment of *Ercc1^-/^*^Δ^ mice was conducted twice per week to evaluate age-related symptoms, including body weight, tremor, forelimb grip strength, kyphosis, hindlimb paralysis, gait disorder, dystonia and ataxia. Kyphosis, body condition and coat condition were used to reflect general health conditions. Ataxia, dystonia, gait disorder and tremor were used as indicators of aging-related neurodegeneration. Muscle wasting was studied by monitoring hindlimb paralysis and forelimb grip strength. All aging symptoms were scored based on a scale of 0, 0.5 and 1, with the exception of dystonia that has a scale from 0 to 5. The sum of aging scores of each group was used to determine the overall aging conditions, with zero means no symptom presented.

### RT-qPCR

Total RNA was extracted from cells or snap frozen tissues using Trizol reagent (Thermo Fisher, USA). cDNA was synthesized using High-Capacity cDNA Reverse Transcription Kit (Thermo Fisher, USA). Quantitative PCR reactions were performed with PowerUp™ SYBR™ Green Master Mix (ThermoFisher, USA). The experiments were performed according to the manufacturer’s instructions. Fold changes in expression were calculated using the ΔΔCT method. The Gapdh gene was used to normalize results. The primer sequences used are listed below.

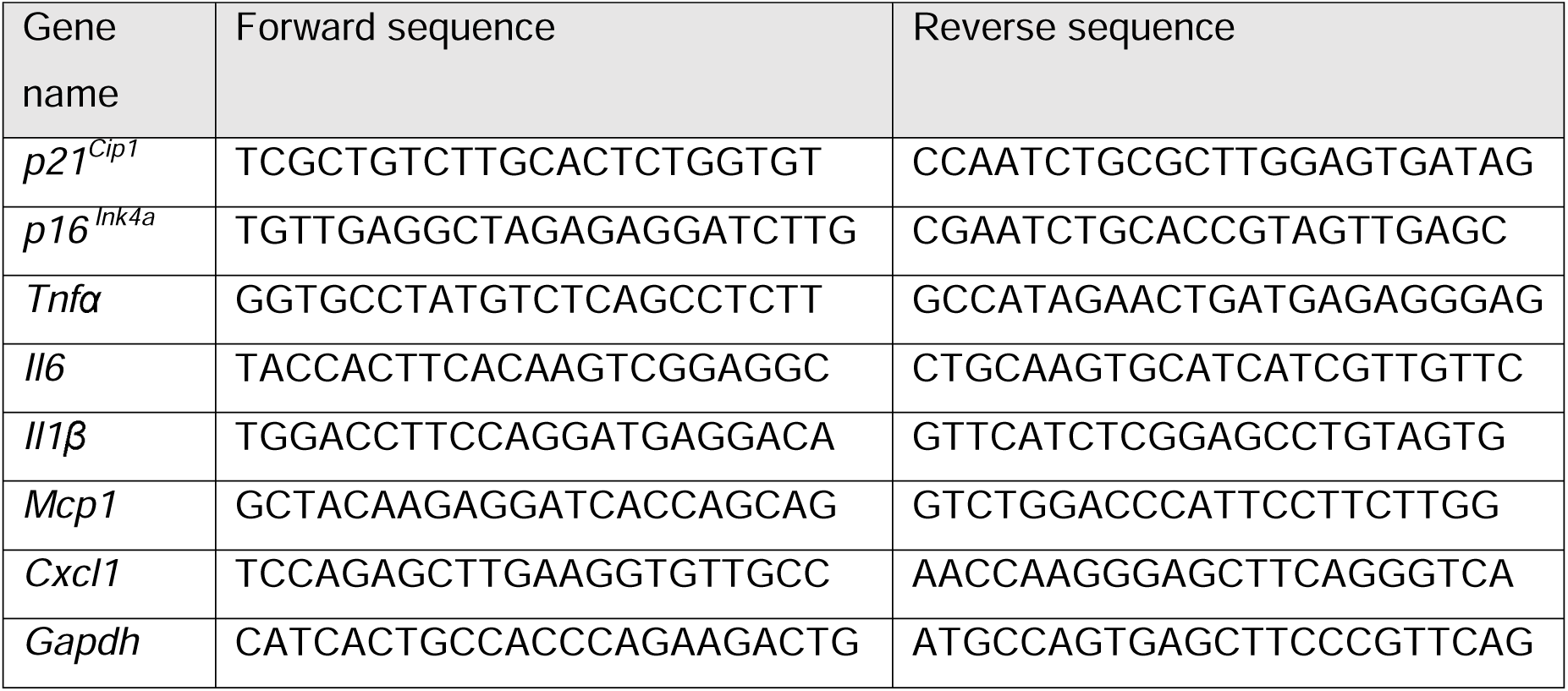

### SIRT6 deacetylation assay

SIRT6 deacetylase activity was assessed using a fluorometric deacetylase assay kit (Sigma-Aldrich, EPI017) according to the manufacturer’s instructions. Assays were conducted in a 96-well black flat-bottom plate, with and without fucoidan treatment. Fluorescence intensity (excitation/emission = 400/505 nm) was measured using a Varioskan™ LUX multimode microplate reader (Thermo Fisher Scientific, USA).

### SIRT6 self-ribosylation assay

SIRT6 self-ribosylation activity was evaluated using a gel-based assay with fluorescein-labeled NAD⁺ (Fluorescein-NAD⁺, Trevigen, 6574). The reaction buffer contained 50 mM Tris-HCl (pH 7.5), 10 µM ZnCl_2_, 150 mM NaCl, and freshly supplemented with 0.5 mM DTT. GST-tagged recombinant human SIRT6 (BioVision, 7697-100) was used at 2 µg per reaction.

Reactions were assembled by preparing a master mix containing all components except the enzyme and Fluorescein-NAD⁺. SIRT6 was added to PCR tubes containing fucoidan or control, followed by addition of the master mix. After gentle mixing, samples were incubated at 37°C for 30 minutes. Fluorescein-NAD⁺ was then added to a final concentration of 25 µM, and the reactions were incubated for an additional 60 minutes at 37°C. Reactions were terminated by adding Laemmli SDS loading buffer supplemented with 10% 2-mercaptoethanol, followed by boiling at 95°C for 5 minutes. Proteins were resolved on 4-20% Tris-glycine SDS-PAGE gels, and fluorescence was imaged using an iBright FL1000 imaging system (Thermo Fisher Scientific) under the 555 nm channel (excitation: 475-490 nm; emission: 510-520 nm). A duplicate gel was stained with Coomassie Brilliant Blue G-250 to confirm equal protein loading.

### siRNA transfection

Transfections were performed using Lipofectamine™ 3000 (Invitrogen) according to the manufacturer’s protocol. Silencer™ Select Pre-Designed siRNA targeting SIRT6 (Thermo Fisher Scientific, Catalog #: 4392420, siRNA ID: s195228) and a Silencer™ Select Negative Control siRNA (Thermo Fisher Scientific, Catalog #: 4390843) were used. Human fibroblast cells containing the NHEJ-GFP reporter, SIRT6 knockout cells, or senescent MEFs were seeded in 96-well black flat-bottom plates at a density to reach 70-90% confluence at the time of transfection. Cells were transfected using Lipofectamine™ 3000 for 6 hours. Twenty-four hours post-transfection, cells were treated with or without fucoidan for 48 hours according to the experimental design.

### RNA-Seq and enrichment analysis

After 12 hours of treatment, all RNA samples were extracted using Trizol reagent (Thermo Fisher, USA). Three independent biological replicates were analyzed per group. Samples were quantified using fluorimetry (RiboGreen assay) and RNA integrity was assessed using capillary electrophoresis. All samples had at least 500 ng mass and RNA Integrity Number (RIN) of at least 8. Library preparation was carried out using the Illumina TruSeq Stranded Total RNA Library Prep kit, followed by sequencing on the NovaSeq 6000 using 150 PE flow cell, with a sequencing depth of 20 million reads per sample. Quality control on raw sequence data for each sample was performed with FastQC (v0.12.1). Read mapping was performed via STAR (v2.7.11b) using the mouse genome (GRCm39.109) as reference. RSEM (v1.3.1) was used for gene quantification. Differentially expressed genes (DEGs) were quantified using the R package DESeq2 (v1.44.0).^51^ DEGs with adjusted p-values < 0.05 and log2FC > |1| were selected for downstream representation analysis. Gene Set Enrichment Analysis (GSEA) and GO enrichment analysis was performed using Clusterprofiler (v4.12.6).^52^ Further analysis and visualization were done using the gsea module from the GSEApy/Enrichr implementation and the seaborn visualization library (Spyder IDE standalone v6.0.5; Python 3.11.11 platform).

For interaction network construction, sets of gene identifiers for DEGs were used as node inputs in the STRING interaction database (https://string-db.org). Results were restricted to M. musculus and the submitted list of genes only. Molecular interaction modes (edges) with highest confidence score (>0.90) were generated and used to analyze highly significant (FDR<0.01) functional enrichment of GO and KEGG terms. Subsequently, nodes and edges were imported to CytoScape (v3.9.0) using the stringApp (v1.7.0) add-on. Subsequently, nodes were visualized using yFiles layout algorithm and clustered using clusterMaker2 (v2.0) MCL clustering with a granularity parameter of 2.5 for all networks.

### Statistical analysis

All data are shown as either mean ± SD or mean ± SEM, unless otherwise indicated. For comparison between two groups, unpaired, two-tailed Student’s t-tests were used. Analysis of variance (ANOVA), followed by a post hoc test for multiple comparisons (Dunnett’s), was used for comparison of groups of three or more. GraphPad Prism software was used for statistical analysis. A value of *p* < 0.05 was considered as statistically significant, shown as \**p* < 0.05, \*\**p* < 0.01, \*\*\**p* < 0.001 and \*\*\*\**p* < 0.0001.

---

**Supplementary Figure S1. Senotherapeutic activity of Fucoidan-FV in various senescent cell models, related to** Figure 1.

Fucoidan-FV reduced senescence detected by staining with C_12_FDG for SA-β-gal activity in (**A**) senescent WT MEFs induced by hydrogen peroxide, (**B**) senescent WT MEFs induced by etoposide, (**C**) senescent IMR90 induced by etoposide, (**D**) senescent HUVECs induced by proliferative passaging. Representative images were captured using Cytation 1 at 4X magnification. Blue fluorescence indicates Hoechst 33324-stained nuclei and green fluorescence marks C_12_FDG-stained SA-β-gal positive senescent cells.

**Supplementary Figure S2. Senotherapeutic activity of Fucoidan-MP in various senescent cell models, related to** Figure 1.

Fucoidan-MP reduced senescence detected by staining with C_12_FDG for SA-β-gal activity in (**A**) senescent WT MEFs induced by hydrogen peroxide, (**B**) senescent WT MEFs induced by etoposide, (**C**) senescent IMR90 induced by etoposide, (**D**) senescent HUVECs induced by proliferative passaging. Representative images were captured using Cytation 1 at 4X magnification. Blue fluorescence indicates Hoechst 33324-stained nuclei and green fluorescence marks C_12_FDG-stained SA-β-gal positive senescent cells.

**Supplementary Figure S3. Senotherapeutic effects of Fucoidan-FV and Fucoidan-MP in aged WT C57BL/6 mice, related to** Figure 2.

(**A**) The effects of Fucoidan-FV on reducing senescence markers across different tissues in WT C57BL/6 mice aged 32 months. Error bars represent SEM for n = 6. (**B**) The effects of Fucoidan-MP on reducing senescence across different tissues in WT C57BL/6 mice aged 32 months. Error bars represent SEM for n = 6.

**Supplementary Figure S4. Transcriptional changes in senescent MEFs and additional RNA-Seq analysis, related to** Figure 4.

(**A**) Principal component analysis plot for RNA-Seq data obtained from untreated (UT) or Fucoidan-FV treated non-senescent (NS/CTRL) and senescent (Sen/Eto) WT MEFs. (**B**) Normalized expression of CDKN1A and CDKN2A in UT NS and UT Sen MEFs; n = 3 per group. (**C**) Volcano plot highlighting differentially expressed SASP genes between UT NS and UT Sen MEF; cutoffs at |log_2_FC|=1 and -log_10_padj = 1.3. (**D**) Venn diagrams for comparisons among different treatment groups. (**E**) Radial interaction network diagram of filtered downregulated DEGs in UT Sen vs 100 µg/mL Fucoidan-FV treated MEFs. Node size corresponds to degree centrality in network topology; edge width corresponds to undirected confidence of interaction between nodes. Top 4 MCL clusters are annotated with colored nodes and labeled with the highest enriched GO term. (**F**) Top 20 GO biological processes (GOBP) pathways overrepresented by filtered up- and downregulated DEGs in UT Sen vs 100 µg/mL Fucoidan-FV treated MEFs.

## REFERENCES

1. Lopez-Otin, C., Blasco, M.A., Partridge, L., Serrano, M., and Kroemer, G. (2013). The hallmarks of aging. Cell 153, 1194–1217.

2. Kennedy, B.K., Berger, S.L., Brunet, A., Campisi, J., Cuervo, A.M., Epel, E.S., Franceschi, C., Lithgow, G.J., Morimoto, R.I., Pessin, J.E., et al. (2014). Geroscience: linking aging to chronic disease. Cell 159, 709–713.

3. López-Otín, C., Blasco, M.A., Partridge, L., Serrano, M., and Kroemer, G. (2023). Hallmarks of aging: An expanding universe. Cell 186, 243–278.

4. Hayflick, L., and Moorhead, P.S. (1961). The serial cultivation of human diploid cell strains. Exp Cell Res 25, 585–621.

5. Hernandez-Segura, A., Nehme, J., and Demaria, M. (2018). Hallmarks of Cellular Senescence. Trends Cell Biol 28, 436–453.

6. Dimri, G.P., Lee, X., Basile, G., Acosta, M., Scott, G., Roskelley, C., Medrano, E.E., Linskens, M., Rubelj, I., Pereira-Smith, O., et al. (1995). A biomarker that identifies senescent human cells in culture and in aging skin in vivo. Proc Natl Acad Sci U S A 92, 9363–9367.

7. Coppe, J.P., Desprez, P.Y., Krtolica, A., and Campisi, J. (2010). The senescence-associated secretory phenotype: the dark side of tumor suppression. Annu Rev Pathol 5, 99–118.

8. Birch, J., and Gil, J. (2020). Senescence and the SASP: many therapeutic avenues. Genes Dev 34, 1565–1576.

9. Gorgoulis, V., Adams, P.D., Alimonti, A., Bennett, D.C., Bischof, O., Bishop, C., Campisi, J., Collado, M., Evangelou, K., Ferbeyre, G., et al. (2019). Cellular Senescence: Defining a Path Forward. Cell 179, 813–827.

10. Wiley, C.D., and Campisi, J. (2021). The metabolic roots of senescence: mechanisms and opportunities for intervention. Nat Metab 3, 1290–1301.

11. Baker, D.J., Wijshake, T., Tchkonia, T., LeBrasseur, N.K., Childs, B.G., van de Sluis, B., Kirkland, J.L., and van Deursen, J.M. (2011). Clearance of p16Ink4a-positive senescent cells delays ageing-associated disorders. Nature 479, 232–236.

12. Baker, D.J., Childs, B.G., Durik, M., Wijers, M.E., Sieben, C.J., Zhong, J., A. Saltness, R., Jeganathan, K.B., Verzosa, G.C., Pezeshki, A., et al. (2016). Naturally occurring p16Ink4a-positive cells shorten healthy lifespan. Nature 530, 184–189.

13. Childs, B.G., Durik, M., Baker, D.J., and van Deursen, J.M. (2015). Cellular senescence in aging and age-related disease: from mechanisms to therapy. Nat Med 21, 1424–1435.

14. He, S., and Sharpless, N.E. (2017). Senescence in Health and Disease. Cell 169, 1000–1011.

15. Wang, B., Wang, L., Gasek, N.S., Kuo, C.L., Nie, J., Kim, T., Yan, P., Zhu, J., Torrance, B.L., Zhou, Y., et al. (2024). Intermittent clearance of p21-highly-expressing cells extends lifespan and confers sustained benefits to health and physical function. Cell Metab 36, 1795–1805 e1796.

16. Zhang, L., Pitcher, L.E., Yousefzadeh, M.J., Niedernhofer, L.J., Robbins, P.D., and Zhu, Y. (2022). Cellular senescence: a key therapeutic target in aging and diseases. J Clin Invest 132, e158450.

17. Prasnikar, E., Borisek, J., and Perdih, A. (2020). Senescent cells as promising targets to tackle age-related diseases. Ageing Res Rev 66, 101251.

18. Zhang, L., Pitcher, L.E., Prahalad, V., Niedernhofer, L.J., and Robbins, P.D. (2023). Targeting cellular senescence with senotherapeutics: senolytics and senomorphics. FEBS J 290, 1362–1383.

19. Zhang, L., Pitcher, L.E., Prahalad, V., Niedernhofer, L.J., and Robbins, P.D. (2021). Recent advances in the discovery of senolytics. Mech Ageing Dev 200, 111587.

20. Borghesan, M., Hoogaars, W.M.H., Varela-Eirin, M., Talma, N., and Demaria, M. (2020). A Senescence-Centric View of Aging: Implications for Longevity and Disease. Trends Cell Biol 30, 777–791.

21. Di Micco, R., Krizhanovsky, V., Baker, D., and d’Adda di Fagagna, F. (2021). Cellular senescence in ageing: from mechanisms to therapeutic opportunities. Nat Rev Mol Cell Biol 22, 75–95.

22. Wang, Y., Xing, M., Cao, Q., Ji, A., Liang, H., and Song, S. (2019). Biological Activities of Fucoidan and the Factors Mediating Its Therapeutic Effects: A Review of Recent Studies. Marine Drugs 17, 183.

23. Oliveira, C., Neves, N.M., Reis, R.L., Martins, A., and Silva, T.H. (2020). A review on fucoidan antitumor strategies: From a biological active agent to a structural component of fucoidan-based systems. Carbohydrate Polymers 239, 116131.

24. Apostolova, E., Lukova, P., Baldzhieva, A., Katsarov, P., Nikolova, M., Iliev, I., Peychev, L., Trica, B., Oancea, F., Delattre, C., et al. (2020). Immunomodulatory and Anti-Inflammatory Effects of Fucoidan: A Review. Polymers (Basel) 12.

25. Luthuli, S., Wu, S., Cheng, Y., Zheng, X., Wu, M., and Tong, H. (2019). Therapeutic Effects of Fucoidan: A Review on Recent Studies. Mar Drugs 17, 487.

26. Jin, J.O., Chauhan, P.S., Arukha, A.P., Chavda, V., Dubey, A., and Yadav, D. (2021). The Therapeutic Potential of the Anticancer Activity of Fucoidan: Current Advances and Hurdles. Mar Drugs 19.

27. Fitton, J.H., Stringer, D.N., Park, A.Y., and Karpiniec, S.S. (2019). Therapies from Fucoidan: New Developments. Marine Drugs 17, 571.

28. Ale, M.T., Mikkelsen, J.D., and Meyer, A.S. (2011). Important Determinants for Fucoidan Bioactivity: A Critical Review of Structure-Function Relations and Extraction Methods for Fucose-Containing Sulfated Polysaccharides from Brown Seaweeds. Marine Drugs 9, 2106–2130.

29. Morya, V.K., Kim, J., and Kim, E.-K. (2012). Algal fucoidan: structural and size-dependent bioactivities and their perspectives. Applied Microbiology and Biotechnology 93, 71–82.

30. Cumashi, A., Ushakova, N.A., Preobrazhenskaya, M.E., D’Incecco, A., Piccoli, A., Totani, L., Tinari, N., Morozevich, G.E., Berman, A.E., Bilan, M.I., et al. (2007). A comparative study of the anti-inflammatory, anticoagulant, antiangiogenic, and antiadhesive activities of nine different fucoidans from brown seaweeds. Glycobiology 17, 541–552.

31. Niedernhofer, L.J., Garinis, G.A., Raams, A., Lalai, A.S., Robinson, A.R., Appeldoorn, E., Odijk, H., Oostendorp, R., Ahmad, A., van Leeuwen, W., et al. (2006). A new progeroid syndrome reveals that genotoxic stress suppresses the somatotroph axis. Nature 444, 1038.

32. Fuhrmann-Stroissnigg, H., Ling, Y.Y., Zhao, J., McGowan, S.J., Zhu, Y., Brooks, R.W., Grassi, D., Gregg, S.Q., Stripay, J.L., Dorronsoro, A., et al. (2017). Identification of HSP90 inhibitors as a novel class of senolytics. Nat Commun 8, 422.

33. Zhang, Y.Z., Naleway, J.J., Larison, K.D., Huang, Z.J., and Haugland, R.P. (1991). Detecting lacZ gene expression in living cells with new lipophilic, fluorogenic beta-galactosidase substrates. Faseb j 5, 3108–3113.

34. Bruno, M.E.C., Mukherjee, S., Powell, W.L., Mori, S.F., Wallace, F.K., Balasuriya, B.K., Su, L.C., Stromberg, A.J., Cohen, D.A., and Starr, M.E. (2022). Accumulation of γδ T cells in visceral fat with aging promotes chronic inflammation. GeroScience 44, 1761–1778.

35. Gurkar, A.U., and Niedernhofer, L.J. (2015). Comparison of mice with accelerated aging caused by distinct mechanisms. Exp Gerontol 68, 43–50.

36. Saul, D., Kosinsky, R.L., Atkinson, E.J., Doolittle, M.L., Zhang, X., LeBrasseur, N.K., Pignolo, R.J., Robbins, P.D., Niedernhofer, L.J., Ikeno, Y., et al. (2022). A new gene set identifies senescent cells and predicts senescence-associated pathways across tissues. Nat Commun 13, 4827.

37. Rahnasto-Rilla, M.K., McLoughlin, P., Kulikowicz, T., Doyle, M., Bohr, V.A., Lahtela-Kakkonen, M., Ferrucci, L., Hayes, M., and Moaddel, R. (2017). The Identification of a SIRT6 Activator from Brown Algae Fucus distichus. Mar Drugs 15.

38. Korotkov, A., Seluanov, A., and Gorbunova, V. (2021). Sirtuin 6: linking longevity with genome and epigenome stability. Trends Cell Biol 31, 994–1006.

39. Guo, Z., Li, P., Ge, J., and Li, H. (2022). SIRT6 in Aging, Metabolism, Inflammation and Cardiovascular Diseases. Aging Dis 13, 1787–1822.

40. Chang, A.R., Ferrer, C.M., and Mostoslavsky, R. (2020). SIRT6, a Mammalian Deacylase with Multitasking Abilities. Physiol Rev 100, 145–169.

41. Michishita, E., McCord, R.A., Boxer, L.D., Barber, M.F., Hong, T., Gozani, O., and Chua, K.F. (2009). Cell cycle-dependent deacetylation of telomeric histone H3 lysine K56 by human SIRT6. Cell Cycle 8, 2664–2666.

42. Michishita, E., McCord, R.A., Berber, E., Kioi, M., Padilla-Nash, H., Damian, M., Cheung, P., Kusumoto, R., Kawahara, T.L.A., Barrett, J.C., et al. (2008). SIRT6 is a histone H3 lysine 9 deacetylase that modulates telomeric chromatin. Nature 452, 492.

43. Mao, Z., Hine, C., Tian, X., Van Meter, M., Au, M., Vaidya, A., Seluanov, A., and Gorbunova, V. (2011). SIRT6 Promotes DNA Repair Under Stress by Activating PARP1. Science 332, 1443–1446.

44. Cao, L., Lee, S.G., Lim, K.T., and Kim, H.-R. (2020). Potential Anti-Aging Substances Derived from Seaweeds. Marine Drugs 18, 564.

45. Salekeen, R., Joydip, B., Rani, S.P., Didarul, I.K.M., Emdadul, I.M., Morsaline, B.M., and and Rahman, S.M.M. (2022). Marine phycocompound screening reveals a potential source of novel senotherapeutics. Journal of Biomolecular Structure and Dynamics 40, 6071–6085.

46. Jackson, M.D., and Denu, J.M. (2002). Structural identification of 2’- and 3’-O-acetyl-ADP-ribose as novel metabolites derived from the Sir2 family of beta -NAD+-dependent histone/protein deacetylases. J Biol Chem 277, 18535–18544.

47. Simon, M., Yang, J., Gigas, J., Earley, E.J., Hillpot, E., Zhang, L., Zagorulya, M., Tombline, G., Gilbert, M., Yuen, S.L., et al. (2022). A rare human centenarian variant of SIRT6 enhances genome stability and interaction with Lamin A. EMBO J 41, e110393.

48. Simon, M., Van Meter, M., Ablaeva, J., Ke, Z., Gonzalez, R.S., Taguchi, T., De Cecco, M., Leonova, K.I., Kogan, V., Helfand, S.L., et al. (2019). LINE1 Derepression in Aged Wild-Type and SIRT6-Deficient Mice Drives Inflammation. Cell Metab 29, 871–885 e875.

49. Imbs, T.I., Zvyagintseva, T.N., and Ermakova, S.P. (2020). Is the transformation of fucoidans in human body possible? International Journal of Biological Macromolecules 142, 778–781.

50. Citkowska, A., Szekalska, M., and Winnicka, K. (2019). Possibilities of Fucoidan Utilization in the Development of Pharmaceutical Dosage Forms. Mar Drugs 17.

51. Love, M.I., Huber, W., and Anders, S. (2014). Moderated estimation of fold change and dispersion for RNA-seq data with DESeq2. Genome Biology 15, 550.

52. Xu, S., Hu, E., Cai, Y., Xie, Z., Luo, X., Zhan, L., Tang, W., Wang, Q., Liu, B., Wang, R., et al. (2024). Using clusterProfiler to characterize multiomics data. Nature Protocols 19, 3292–3320.

