## Supplementary figures and images for "Fucoidans are senotherapeutics that enhance SIRT6-dependent DNA repair"

### Supplementary Figure S1

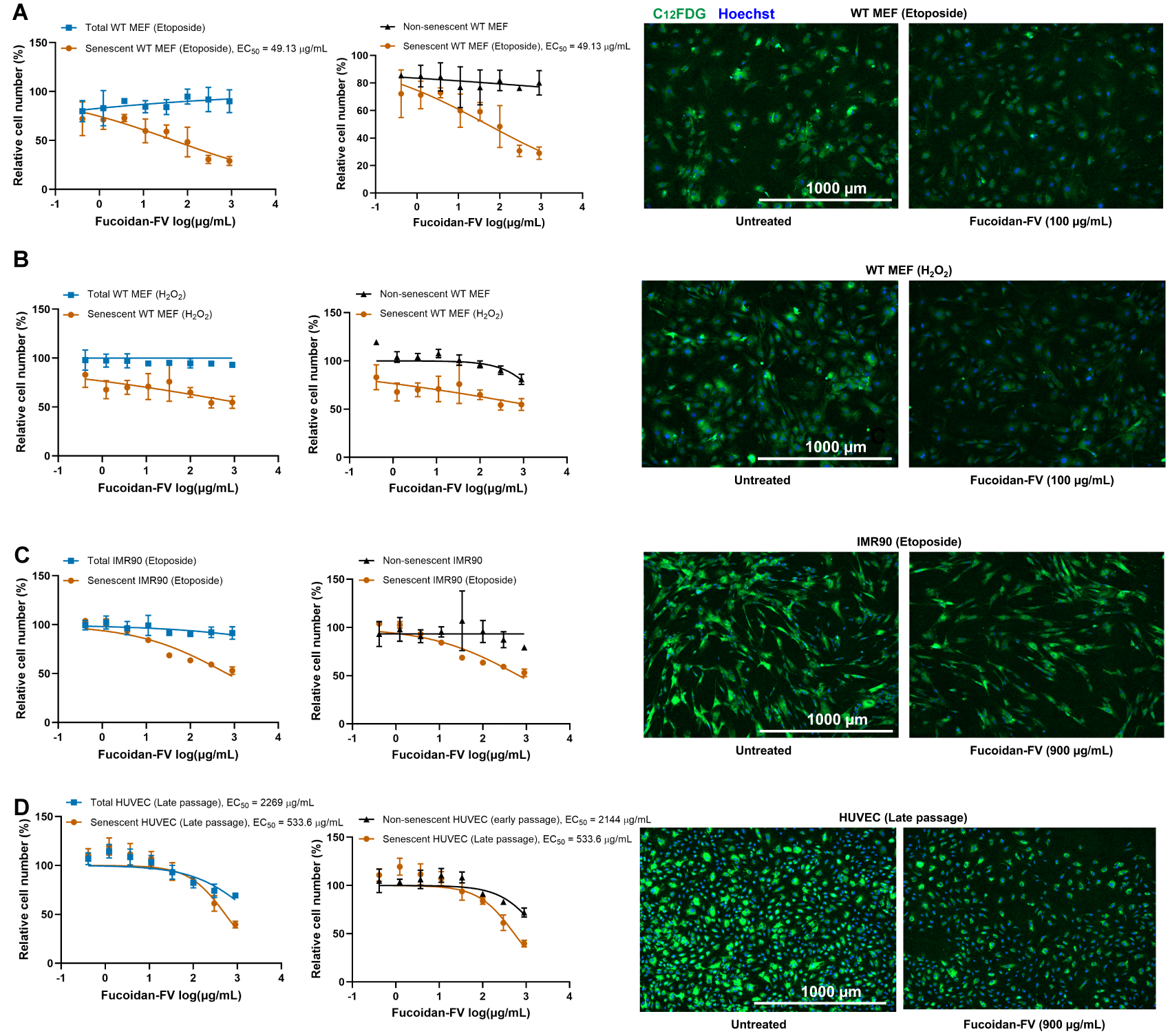

### Supplementary Figure S2

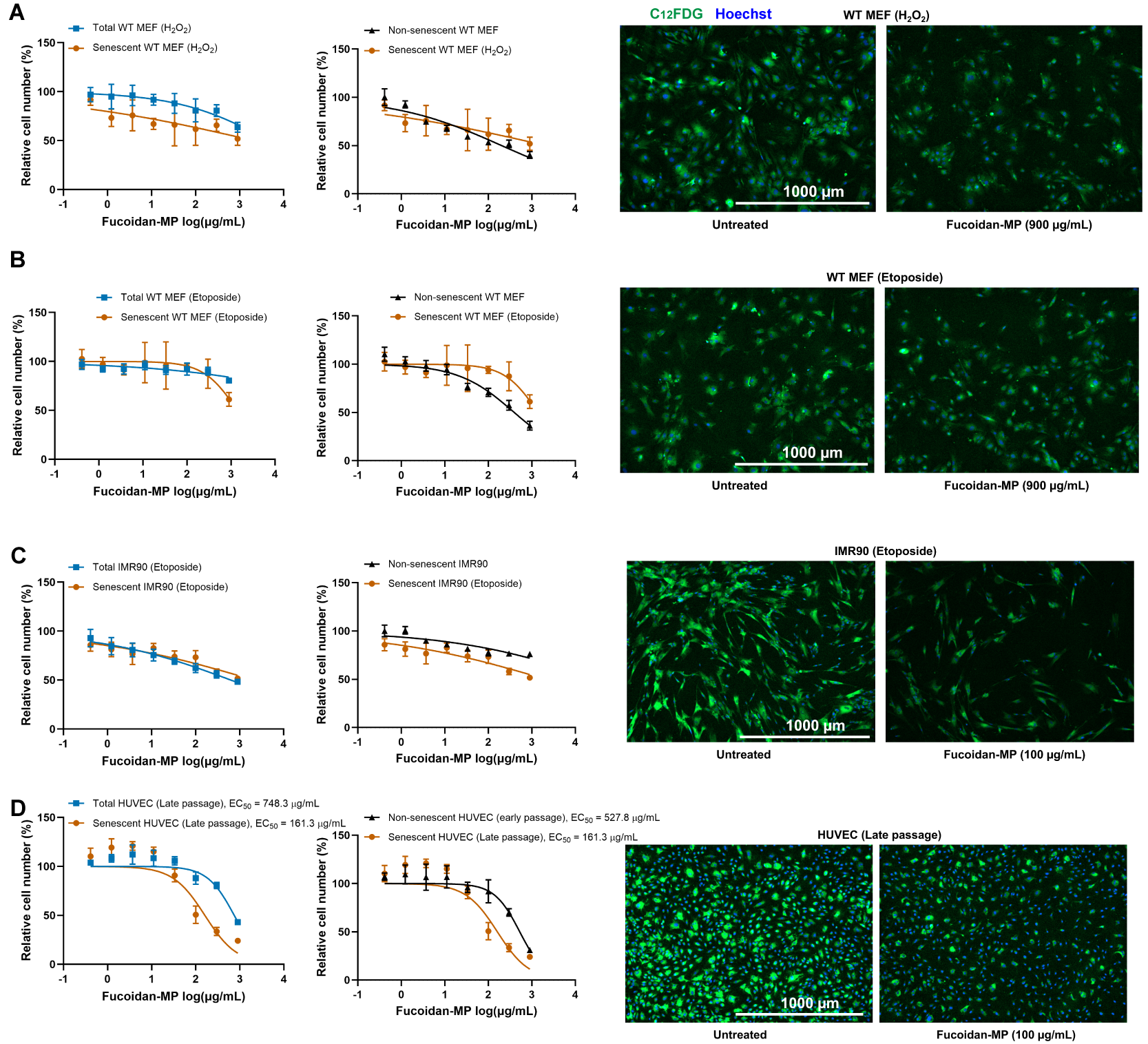

### Supplementary Figure S3

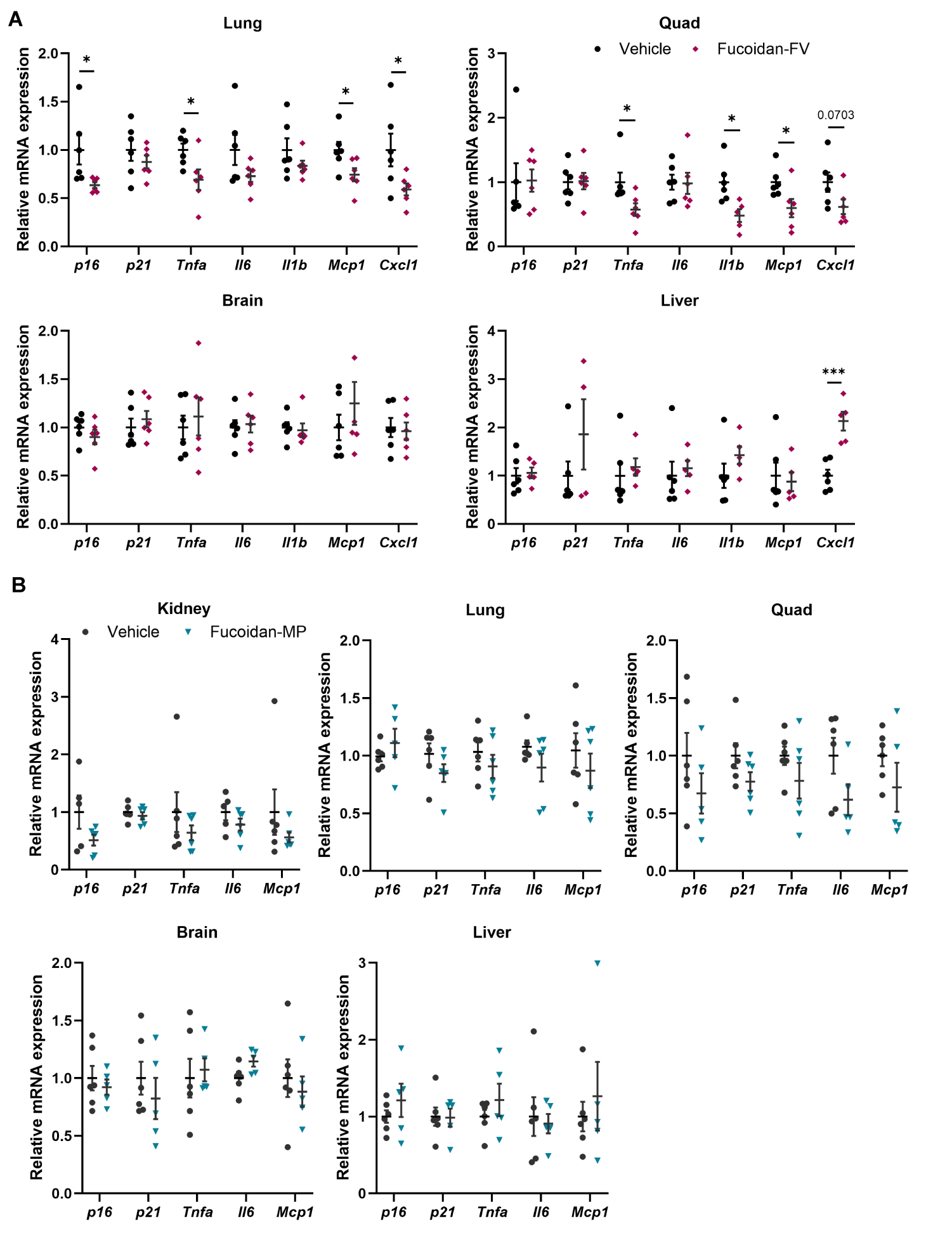

### Supplementary Figure S4

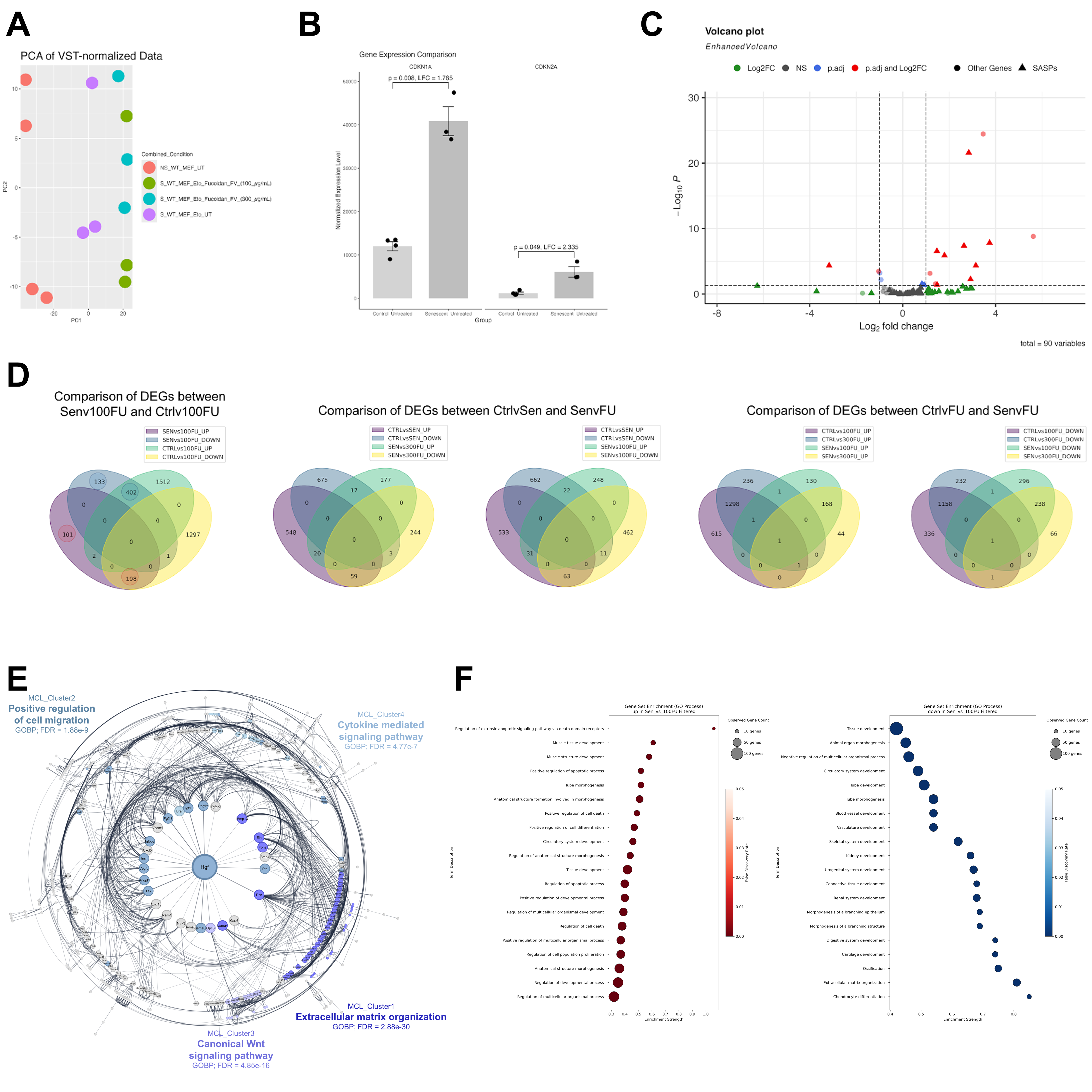
